# Optimization of host cell-compatible, antimicrobial peptides effective against biofilms and clinical isolates of drug-resistant bacteria

**DOI:** 10.1101/2022.12.01.518642

**Authors:** Jenisha Ghimire, Robert J. Hart, Anabel Soldano, Charles H. Chen, Shantanu Guha, Joseph P. Hoffmann, Kalen M. Hall, Leisheng Sun, Benjamin J. Nelson, Timothy K. Lu, Jay K. Kolls, Mario Rivera, Lisa A. Morici, William C. Wimley

## Abstract

Here, we describe the continued synthetic molecular evolution of a lineage of host-compatible antimicrobial peptides (AMP) intended for the treatment of wounds infected with drug-resistant, biofilm-forming bacteria. The peptides tested are variants of an evolved AMP called D-CONGA, which has excellent antimicrobial activities *in vitro* and *in vivo*. In this newest generation of rational D-CONGA variants, we tested multiple sequence-structure-function hypotheses that had not been tested in previous generations. Many of the peptide variants have lower antibacterial activity against Gram-positive or Gram-negative pathogens, especially variants that have altered hydrophobicity, secondary structure potential, or spatial distribution of charged and hydrophobic residues. Thus, D-CONGA is generally well tuned for antimicrobial activity. However, we identified a variant, D-CONGA-Q7, with a polar glutamine inserted into the middle of the sequence, that has higher activity against both planktonic and biofilm-forming bacteria as well as lower cytotoxicity against human fibroblasts. Against clinical isolates of *K. pneumoniae*, innate resistance to D-CONGA was surprisingly common despite a lack of inducible resistance in *P. aeruginosa* reported previously. Yet, these same isolates were susceptible to D-CONGA-Q7. D-CONGA-Q7 is much less vulnerable to AMP resistance in Gram-negative bacteria than its predecessor. Consistent with the spirit of synthetic molecular evolution, D-CONGA-Q7 achieved a critical gain-of-function and has a significantly better activity profile.

## Introduction

Membrane permeabilizing antimicrobial peptides have long been recognized as a promising, but mostly unfulfilled, chemotype in the development of new drugs to treat drug-resistant bacterial infections^1–3^. Thousands of AMPs have been described in the literature^4^, many with potent, broad-spectrum antimicrobial activity against bacterial pathogens *in vitro*. However, due to multiple known impediments to AMP activity *in vivo*, ^5–7^ few have reached late-stage clinical trials. Only polymyxin (colistin) and daptomycin, both naturally occurring lipopeptides, have been approved for use in humans in the US and Europe.

An AMP that is useful, for example in wound treatment, must simultaneously be optimized for antimicrobial activity against multiple microbes but also for a lack of impediments, including host cell, tissue, and protein binding, as well as cytotoxicity, proteolytic degradation, and low solubility. These various activities are complex, multifactorial, and interdependent such that simultaneous rational optimization is not possible. AMP activity against bacteria, considered alone, is so complex and poorly understood that it defies easy explanation, except in the very broadest terms. It depends, ultimately, on cytoplasmic membrane disruption^1–2^, which requires “interfacial activity”^8^. This property is contingent upon strong membrane binding coupled with imperfect amphipathicity, which disrupts lipid organization in bilayers leading to permeabilization. However, accurately predicting which sequences will be effective at bacterial membrane permeabilization or foreseeing how to improve activity or reduce impediments has never been possible.

As a consequence, most AMP discoveries in the literature are the result of trial and error, based on known sequences or physical-chemical hypotheses. We have embraced the spirit of trial and error to efficiently identify and optimize new “host-compatible” AMPs, lacking the known impediments by using synthetic molecular evolution (SME)^5^. In our work, SME is the orthogonal screening of multiple, iterative generations of small libraries under conditions of increasing clinical relevance. SME enables us to test thousands of closely related sequences to identify AMPs with the best broad-spectrum antimicrobial activity while also down-selecting for known impediments. Our libraries of 5-30,000 members are each designed to test the contribution of a small number of hypothesis-based rational variations. After a generation of screening to identify gain-of-function variants, we have found it useful to carry out rounds of semi-rational variation to test the importance of hypotheses that had not been tested previously in libraries. Such cycles of library screening, followed by rational trial and error, have led to continuous improvements in AMP properties and also to an improved understanding of sequence-function relationships, which can result in more intelligent, next-generation library design.

We have carried out multiple generations of AMP evolution, with improvements at every generation. See **Fig. 1** for a history of this AMP lineage. In the first generation, we screened a *de novo* designed peptide library for members that permeabilize bacteria-like lipid vesicles^9–10^. In parallel, we screened the same library for members with broad spectrum antibacterial activity in simple, defined media^11^. These two screens enabled the identification of two distinct families of potent, broad-spectrum AMPs from a single library. We subsequently showed that these first-generation AMPs, like most others, have a list of impediments to *in vivo* activity: i) they lose activity in the presence of concentrated host cells, ii) they have some residual toxicity^11^, iii) they are rapidly degraded by serum endopeptidases^6^, and iv) they have relatively low solubility. We thus designed and synthesized an iterative second-generation library that was screened for multi-organism, sterilizing activity <u>in the presence of concentrated human erythrocytes</u>^5^, to downselect against host cell and protein binding. We also downselected against cytotoxicity by measuring hemolysis in parallel. We downselected against insoluble peptides by pre-incubating in saline solution prior to assays. Having shown that L- and D-amino acid enantiomers of these AMPs have identical activities^5, 7, 11^, we tested the hits using protease-resistant D-amino acid versions to eliminate proteolytic degradation by serum exopeptidases. This second-generation screen led to the discovery of the so-called D-DBS (D-amino acid <u>D</u>ouble <u>B</u>roth <u>S</u>terilization) peptides, which have broad spectrum sterilizing activity against a panel of Gram-positive and Gram-negative pathogens^5^, **Fig. 1**. Importantly, the DBS peptides are fully active in the presence of concentrated host cells and have high solubility (>1 mM) in physiological saline solution. We created a consensus sequence and subsequently obtained some improvement by the removal of two invariant glycines^5^. This lead peptide D-CONGA (<u>D</u>-amino acid <u>CON</u>sensus with <u>G</u>lycine <u>A</u>bsent), which has been described in detail^5^, is highly soluble, highly stable, and has moderately low cytotoxicity, while also having broad-spectrum, antibacterial, and anti-biofilm activity *in vitro* and *in vivo*, even in the presence of host cells, serum proteins, and tissue^5^. D-CONGA also has the important property of resistance avoidance^5^.

**Figure 1.**
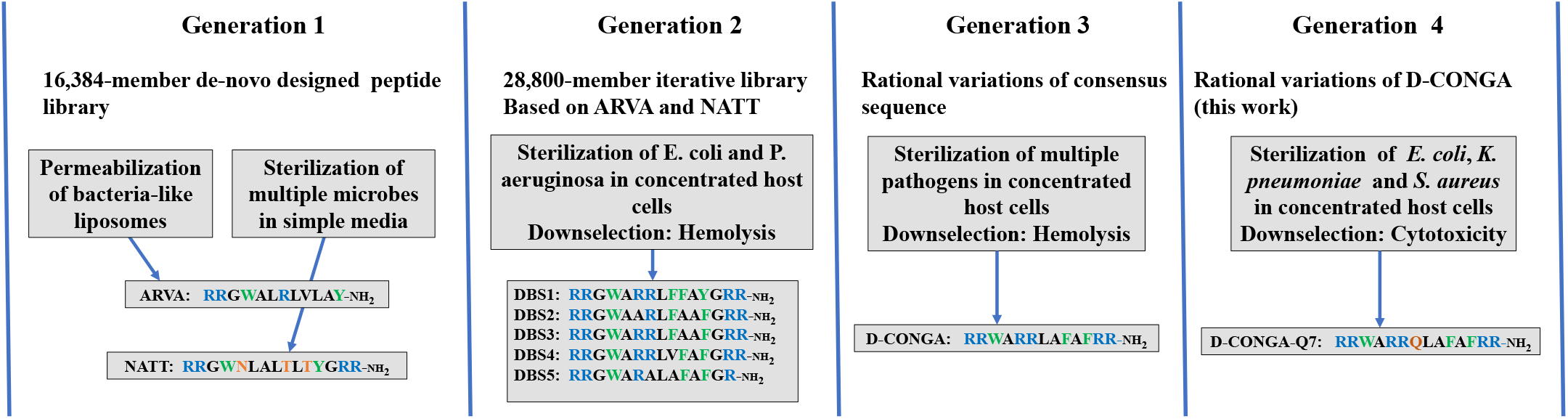
History of the evolution of the AMP lineage discussed in this work. The first generation *de novo* library was screened for liposome permeabilization^9–10^ and for antimicrobial activity in simple media^11^. The two families of AMPs derived from generation 1 were used to design a library, which was screened for host-compatible AMP activity in generation 2^5^. In generation 3, a round of rational variants was designed and tested to identify D-CONGA, a host compatible AMP with highly promising activities *in vitro* and *in vivo*^5^. Generation 4 is the work described here.

In the current work, we subjected the lead peptide D-CONGA to a broad series of rational variations to catalog the important characteristics for its activity as a host-compatible antibiotic. We find that the properties of D-CONGA are mostly well-tuned. Most variations to its sequence or structure propensity either decrease antimicrobial activity and toxicity or have little effect. However, we identified a new variant, called D-CONGA-Q7, with a polar glutamine inserted between the cationic and hydrophobic segments of the peptide, that has significantly improved antibiotic activity, significantly better anti-biofilm activity, and lower residual toxicity compared to D-CONGA. Perhaps most importantly, the newly discovered peptide, D-CONGA-Q7 has significantly improved activity against clinical isolates of drug-resistant bacteria, including isolates of the problematic Gram-negative species *K. pneumoniae* that are resistant to the parent peptide, D-CONGA.

## Results

### Variants of D-CONGA

We previously described the identification of the broad-spectrum, host-cell compatible antimicrobial peptide D-CONGA (**rrwarrlafafrr-amide**) by synthetic molecular evolution^5^. A “host-cell compatible” AMP is defined as one with antibacterial activity that is not inhibited by the presence of concentrated human erythrocytes^5, 7^, which mimics *in vivo* conditions rich with host cells, protein, and tissue. Host cell compatibility is a rare property among known AMPs^5, 7^ but is important to identify in AMPs designed to treat drug-resistant bacterial infections in wounds.

The iterative library from which D-CONGA was derived contained 28,800 members with some variation in the termini and in the core of the peptide, enabling tests of a number of specific hypotheses, described in detail previously^5^. In this work, we created a new set of rational variants to test new hypotheses for improving the activity or reducing the impediments to activity that were not included in the screen and to inform the next generation libraries. The 13 amino acid sequence of D-CONGA (**rrwarrlafafrr-NH2**), shown in **Fig. 2**, consists of critical double arginines on both termini and a double arginine at positions 5 and 6. This architecture creates a highly polar, but also amphipathic, N-terminal hexapeptide sequence of **rrwarr** which has five positive charges, including the N-terminus, plus the aromatic tryptophan. The C-terminal heptapeptide, **lafafrr**, has five consecutive nonpolar residues, including two large, aromatic phenylalanine residues, making for a very hydrophobic, but also amphipathic, segment. We previously showed decreasing hydrophobicity of this segment caused decreased antimicrobial activity, while increasing hydrophobicity caused increased cell toxicity^5^.

**Figure 2.**
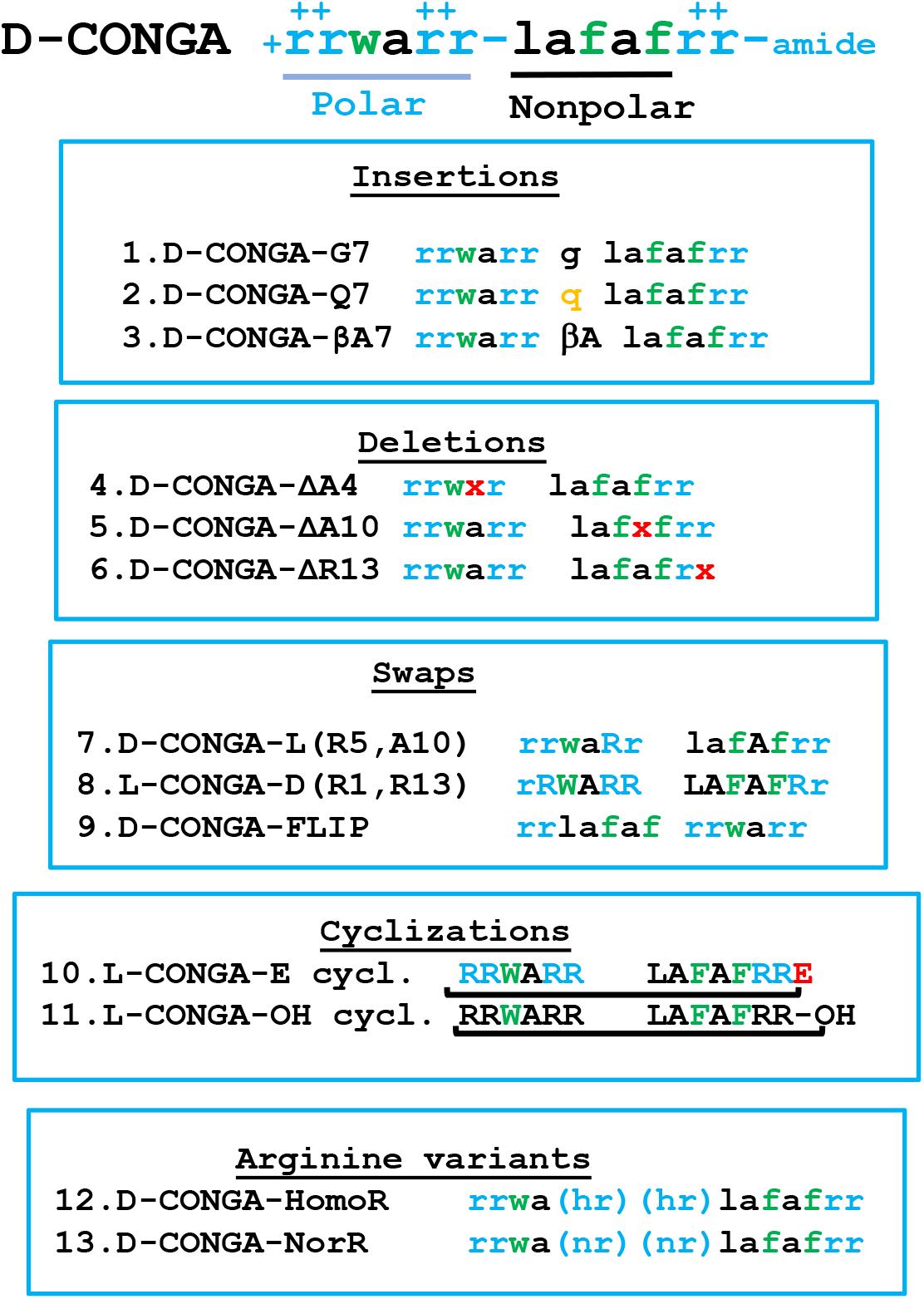
Design and synthesis of D-CONGA variants. 13 rational variants of D-CONGA were synthesized and tested here. The variants can be divided in to five broad groups based on the modifications in the sequence of D-CONGA. <u>Insertions</u> were made by adding an amino acid to the D-CONGA sequence after the R residue at position 6. This position is denoted by a hyphen in the D-CONGA sequence above. <u>Deletions</u> were made by removing an amino acid from D-CONGA that had not been previously varied. <u>Swaps</u> contain 3 variants with position and enantiomer exchanges within CONGA. Cyclized peptides are two variants made by the cyclization of D-CONGA. <u>Arginine variants</u> replace the RR motif at positions 5 and 6 with arginine analogs; norarginine, with one additional methylene unit, and homoargoinine with one fewer methylene unit.

In the molecular evolution and optimization described here, we tested five classes of sequence variations of D-CONGA that had not been previously tested in a library or by rational variation. These are shown in **Fig. 2**. First, we made insertions at position 7 between the independent polar N-terminal and nonpolar C-terminal segments. Glycine and β-alanine were added to increase flexibility between segments, with β-alanine enabling maximum flexibility. These insertions were designed to test the hypothesis that reduced secondary structure propensity can lead to reduced cytotoxicity. We also inserted glutamine at position 7 to test the effect of extending the polar segment by one residue without adding any charges. We chose glutamine for this test because the amide sidechain is the most polar of the unionizable natural amino acids. <u>Second</u>, we tested the effect of deleting alanine residues at positions 4 and 10, because these two alanines had not been varied in any previous generation. These changes enabled the testing of shorter peptides, which are advantageous in a peptide drug candidate. Further, alanine residues do not contribute to polarity or to hydrophobicity^12–13^. In fact, alanine residues reduce amphipathicity wherever they occur. We also deleted the C-terminal arginine because DBS5 was found to be active without it^5^. <u>Third</u>, we switched the chirality of two amino acids within the sequence to assess the effect of interrupting secondary structure, which we hypothesize will reduce cytotoxicity. One such variant peptide had L-amino acids at positions 5 and 10 in the context of an otherwise D-amino acid sequence. These internal positions will maximally interrupt secondary structure. The second such peptide had D-Arg residues on the N- and C-termini of an otherwise L-amino acid peptide. In this peptide, we are also testing the possibility of engineering protease resistance by terminal modification^14^ without using D-amino acids for the whole peptide. In addition to terminal modification, we also made a peptide in which we swapped the N- and C-terminal halves to test for additivity. <u>Fourth</u>, we made cyclized versions of L-CONGA by coupling the N-terminal amino group to either the C-terminus of an L-CONGA peptide acid, or to the sidechain of a C-terminal Asp residue on an L-CONGA peptide-amide. Cyclization is an alternate way to reduce proteolysis of an L-amino acid peptide. It also tests the hypothesis that AMP activity is dependent on a β-sheet or β-hairpin-like secondary structure, which will be increased by cyclization. Fifth, we tested peptides in which the central Arg residues at positions 5 and 6 were replaced with Arg variants with longer or shorter versions of the guanidine-containing sidechains, homo-arginine and nor-arginine, respectively. In this case we are testing a structural hypothesis that the Arg sidechains interact specifically with an anionic component of the bacterial membrane and that the altered length of norR and homoR will alter this interaction.

All peptide variants were synthesized and purified as described previously^5, 7^. They were tested in broth dilution in the presence and absence of 1×10^9^ human RBCs per mL, 20% of the concentration in human blood, against *Escherichia coli, Klebsiella pneumoniae* and *Staphylococcus aureus*. See **Fig. 3A** for MIC values of the variants, compared to D-CONGA which is shown in the top row. *K. pneumoniae* and *S. aureus* were tested here because of their relatively lower sensitivity to D-CONGA. The toxicity of the variants was tested by measuring hemolysis as well as cytotoxicity against human WI-38 fibroblast cells.

**Figure 3.**
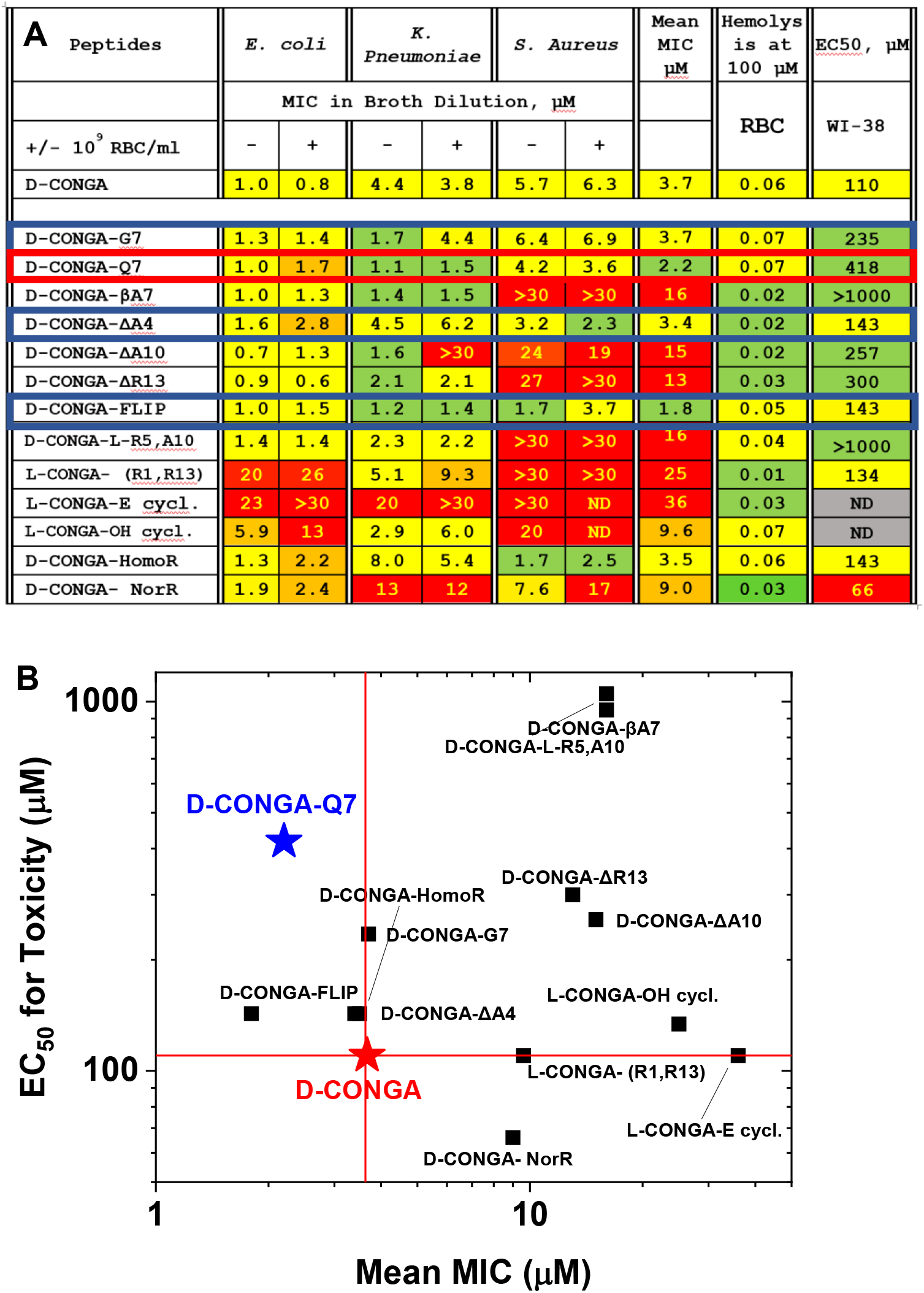
Characterization of D-CONGA variants. **A.** MIC values are reported in μM peptide against gram-negative *E. coli* (EC), *K. pneumoniae* (KP), and the gram-positive pathogen *S. aureus* (SA). The two columns under each organism are for assays performed in the absence (-) and presence (+) of 1×10^9^ human RBC/mL. MIC color codes are as follows: Green: Values are at least 2-fold better than D-CONGA. (Better is defined as lower MIC or higher EC_50_ for cytotoxicity). Yellow: Values are similar to that of D-CONGA (within a factor of two). Orange: Values are more than 2-4-fold worse than D-CONGA. Red: Values are more than 4-fold worse than D-CONGA. “>30” means that sterilization was not observed at 30 μM, the highest concentration tested. The column marked “hemolysis” is the fractional hemolysis of 1×10^8^ human RBCs/mL at 100 μM peptide determined from measurements of serially diluted peptide, starting from 100 μM. The column marked “EC_50_” contains the concentration of peptide that kills 50% of WI-38 human fibroblast cells assayed by entry of SYTOX Green, a DNA binding dye, extrapolated from the experimentally measured range of 0-200 μM. “>1000” signifies that no cytotoxicity was observed at the highest peptide concentration. **B.** A comparison of MIC values and EC_50_ for cytotoxicity divided into four quadrants by the values for D-CONGA. Statistical methods are described in Methods. Both MIC and EC_50_ values have consistent standard errors equal to about 20% of the value of the mean.

### Identification of D-CONGA-Q7, a significantly improved variant

Interestingly, the three bacterial species had different susceptibilities to changes in the sequence of D-CONGA. *E. coli*, against which D-CONGA was highly active, was the least sensitive to changes. The majority of variants have MIC values that are within a factor of two of the MIC for D-CONGA. *K. pneumoniae* had a wider range of changes. Three of the variants were significantly better than D-CONGA, and four of them were significantly worse. *S. aureus* showed the most sensitivity to sequence changes, a phenomenon that we have reported previously^5^. Half of the variants lost all activity against this organism, and only two variants, D-CONGA-Q7 and D-CONGA-FLIP, were better than D-CONGA against *S. aureus*.

Overall, eight of the 13 variants had poorer average antibacterial activity than D-CONGA, **Fig. 3.** Seven of these lost useful activity against *S. aureus* and a few also lost activity against *K. pneumoniae* and *E. coli*. Cyclization had the largest detrimental effect on activity, with both cyclic variants being almost inactive. Deletion of A10 or R13 as well as swapping chirality of two amino acids led to poorer activity, including loss of activity against *S. aureus*. The D-CONGA variants ΔA4, G7 and homoR had roughly the same average MIC as D-CONGA. Only two variants had overall improved antibacterial activity, the insertion variant D-CONGA-Q7 and the flipped D-CONGA-FLIP.

The variant with norarginine was the most cytotoxic variant against fibroblasts and was the only one that was more cytotoxic than the parent, D-CONGA. Four other variants had toxicity against fibroblasts that were similar to D-CONGA and six had lower toxicity. The least toxic variants were the βA7 insertion and the D-CONGA with L-R5 and L-A10, which showed no detectable toxicity, even when extrapolated to 1 mM peptide.

Toxicity values are plotted against average MIC values in **Fig. 3B** which is divided into four quadrants. Only two of the 13 variants of D-CONGA had substantially better activity against all three bacterial pathogens, D-CONGA-Q7 and D-CONGA-FLIP. Of these two, only D-CONGA-Q7 also had lower cytotoxicity against fibroblast cells. Therefore, **we selected D-CONGA-Q7 as the lead peptide** for additional studies against clinical isolates, against biofilms, and in our murine model of infected wounds^5, 15^. In the next sections, we carefully compare these important activities of D-CONGA and D-CONGA-Q7, to demonstrate that D-CONGA-Q7 displays a significant gain-of-function over D-CONGA.

### Activity against clinical isolates of Gram-negative pathogens

Against laboratory strains of bacterial pathogens, D-CONGA-Q7 is measurably better than D-CONGA, **Fig. 3**. However, a much bigger gain-of-function was observed when we compared their activities against clinical isolates of drug-resistant bacteria, with a focus on *K. pneumoniae*, a species of Gram-negative bacteria of rising concern that can show innate resistance to AMPs^16–18^. Two independent sets of isolates were tested, as described in Methods, below. In **Fig. 4** we show the minimum inhibitory concentration (MIC) of eight conventional antibiotics and the two peptides, D-CONGA and D-CONGA-Q7, against the first set of isolates. See **Table 1**. The MIC values, shown in **Fig. 4A,** demonstrate the degree of drug resistance. All isolates are completely resistant to ceftazidime and ampicillin (MIC>150 μM), and 4 of the 14 isolates are resistant to all eight conventional antibiotics tested. D-CONGA, which has excellent activity (MIC ≤ 10 μM) against laboratory strains of all ESKAPE^19^ pathogens, including other strains of *K. pneumoniae*, has poor MIC ≥ 20 μM against 5 of the 14 isolates (36%). Two isolates of *K. pneumoniae* are resistant to D-CONGA as well as to all conventional antibiotics tested. In sharp contrast, D-CONGA-Q7 has substantial activity against all isolates. MIC values for D-CONGA-Q7 are lower than for D-CONGA for all 14 isolates. In **Fig. 4B** we plot the fraction of isolates sterilized as functions of antibiotic concentration. The extraordinary activity of the new variant D-CONGA-Q7 is shown by the fact that half of these clinical isolates are sterilized by 2 μM D-CONGA-Q7, and 12/14 (86%) are sterilized by 5 μM. In comparison, the best conventional antibiotic, ciprofloxacin, is active against only 8 of the 14 isolates, or 57% at 100 μM. To verify these results, we tested D-CONGA and D-CONGA-Q7 against a second, completely independent set of previously described clinical isolates of *K. pneumoniae^20–21^*. The results, in **Fig. 5A,** show the same behavior as above. For *K. pneumoniae* isolates, 7/19 (37%) are resistant to D-CONGA up to 25 μM, and only 2/19 (11%) have MIC < 5 μM. Validating our observations with the first set of isolates, the activity of D-CONGA-Q7 against these isolates is much better. Only 1/21 (5%) of strains are resistant to D-CONGA-Q7, and 9/21 (57%) have MIC < 5 μM. See **Fig. 5B**.

**Table 1.** Peptides and conventional antibiotics. The two antimicrobial peptides, D-CONGA and D-CONGA-Q7, and eight conventional antibiotics from four different drug classes that were evaluated against drug resistant clinical isolates of *K. pneumoniae, P. aeruginosa*, *Acinetobacter baumanii*, and *S. aureus*. Abbreviations for antibiotics used in **Fig. 4** are shown here.

| Compound | Abbr. | Class/Target |
| --- | --- | --- |
| D-CONGA | -- | AMP |
| D-CONGA-Q7 | -- | AMP |
| Gentamycin | GEN | Aminoglycoside |
| Ceftazidime | CAZ | Cephalosporin |
| Ampicillin | AMP | Beta-lactam |
| Ciprofloxacin | CIPRO | Fluoroquinolone |
| Tobramycin | TOB | Aminoglycoside |
| Trimethoprim | TMP | DHFR Inhibitor |
| Ceftriaxone | CRO | Cephalosporin |
| Meropenem | MEM | Carbopenem |

**Figure 4.**
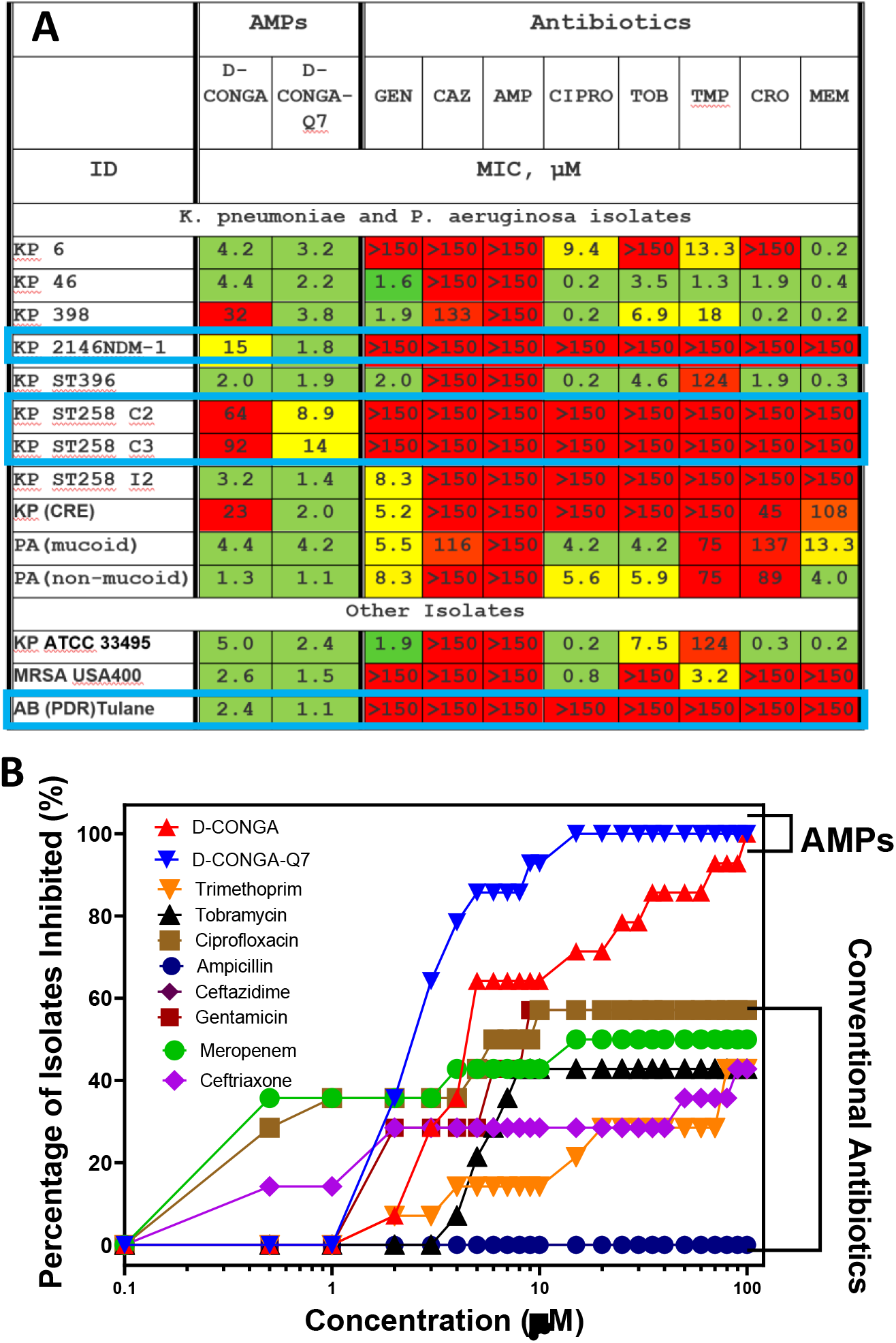
Activity of D-CONGA, D-CONGA-Q7, and conventional antibiotics against clinical isolates. **A.** MIC values in broth dilution are reported in μM concentrations against 14 clinical isolates of resistant bacterial strains. Color codes are as follows: Green: MIC ≤ 5 μM. Yellow: 5≤ MIC ≤ 20. Red: MIC ≥ 20 μM. “>150” means that sterilization was not observed at 150 μM, the highest concentration of antibiotic tested. **B.** Fraction of the 14 isolates sterilized versus antibiotic concentration. Statistical analyses are described below in Methods. MIC values represent at least 3 independent measurements and have consistent standard errors equal to about 40% of the value of the mean.

**Figure 5.**
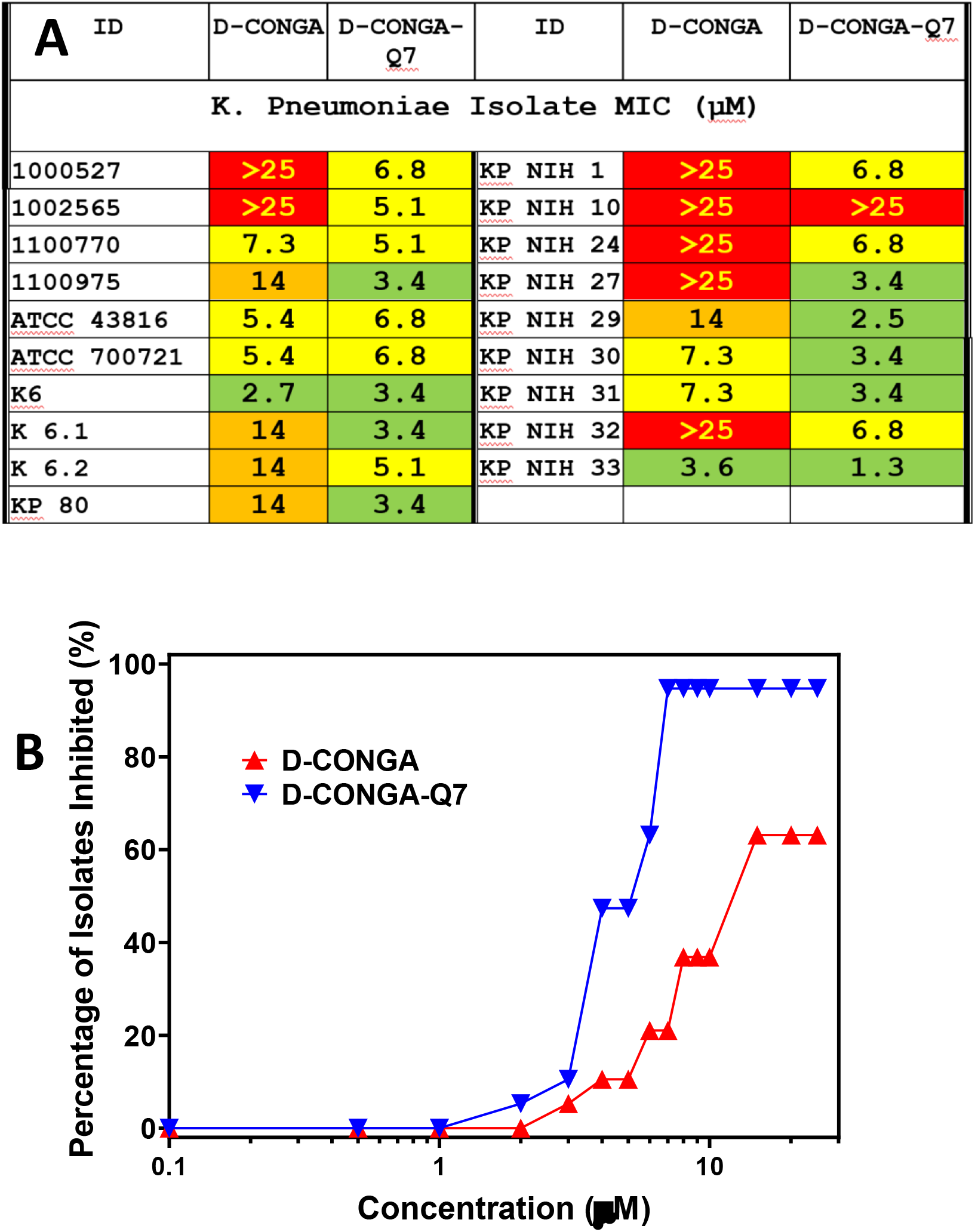
Activity of D-CONGA and D-CONGA-Q7 against an independent set of clinical isolates. **A.** Activity of D-CONGA and D-CONGA-Q7 against an independent set of clinical isolates of *K. pneumoniae* described elsewhere^20–21^. MIC values are shown in **A**. Green cells indicate MIC ≤ 5 μM. Yellow cells indicate MIC ≤ 8 μM. Red cells indicate MIC ≥ 10 μM. **B.** Fraction of the isolates sterilized versus antibiotic concentration. Statistical analyses are described below in Methods. MIC values represent at least 3 independent measurements and have consistent standard errors equal to about 40% of the value of the mean.

#### Antibiofilm activity of D-CONGA-Q7

We reported previously^5^ that D-CONGA has significant activity against biofilms formed by Gram-negative *P. aeruginosa* and Gram-positive *Streptococcus mutans*. Here, we directly compare the antibiofilm activity of D-CONGA and D-CONGA-Q7 using a mature pellicle biofilm of *P. aeruginosa*^22^. Pellicle biofilms are amenable to imaging by confocal laser fluorescence microscopy (CLFM) and to harvesting, which can be desirable for additional biofilm analysis, including counting of viable cells^22–23^. To determine the susceptibility of pellicle biofilms to D-CONGA and D-CONGA-Q7, we cultured 2-day old biofilms as described in Methods. The coverslip-adhered biofilms were exposed to challenge solution for 24 h prior to staining with SYTOX red and imaging with CLFM for GFP, which indicates live cells, and for SYTOX Red which stains dead cells and extracellular DNA. The images obtained from biofilms treated with D-CONGA or D-CONGA-Q7, **Fig. 6A,** show viable cells (yellow) in the untreated control and a concentration-dependent decrease in viable cells, concomitant with an increase in dead cells and extracellular DNA (red), when biofilms are treated with each of the peptides. The % cell survival was quantitated in two ways: (i) COMSTAT was used to compare biofilm biomass in CLFM images by estimating the biofilm biovolume, which is calculated as the overall volume/substratum area (μm^3^/μm^2^)^24–25^. Hence, % cell survival in each of the conditions is expressed as the ratio (untreated biomass)/(treated biomass). (ii) The biofilms were dispersed mechanically for subsequent enumeration of viable cells, such that % survival is expressed as the ratio (CFUuntreated biofilm/CFUtreated biofilm). The results obtained using COMSTAT and viable cell enumeration are shown in Figs. 6B and 6C, respectively. Both assays validate the previous reports of potent antibiofilm activity by D-CONGA. More importantly for this work, the data show that D-CONGA-Q7 is consistently more potent than D-CONGA, reducing viable biofilm bacteria by ≥90% at 8 μM peptide.

**Figure 6.**
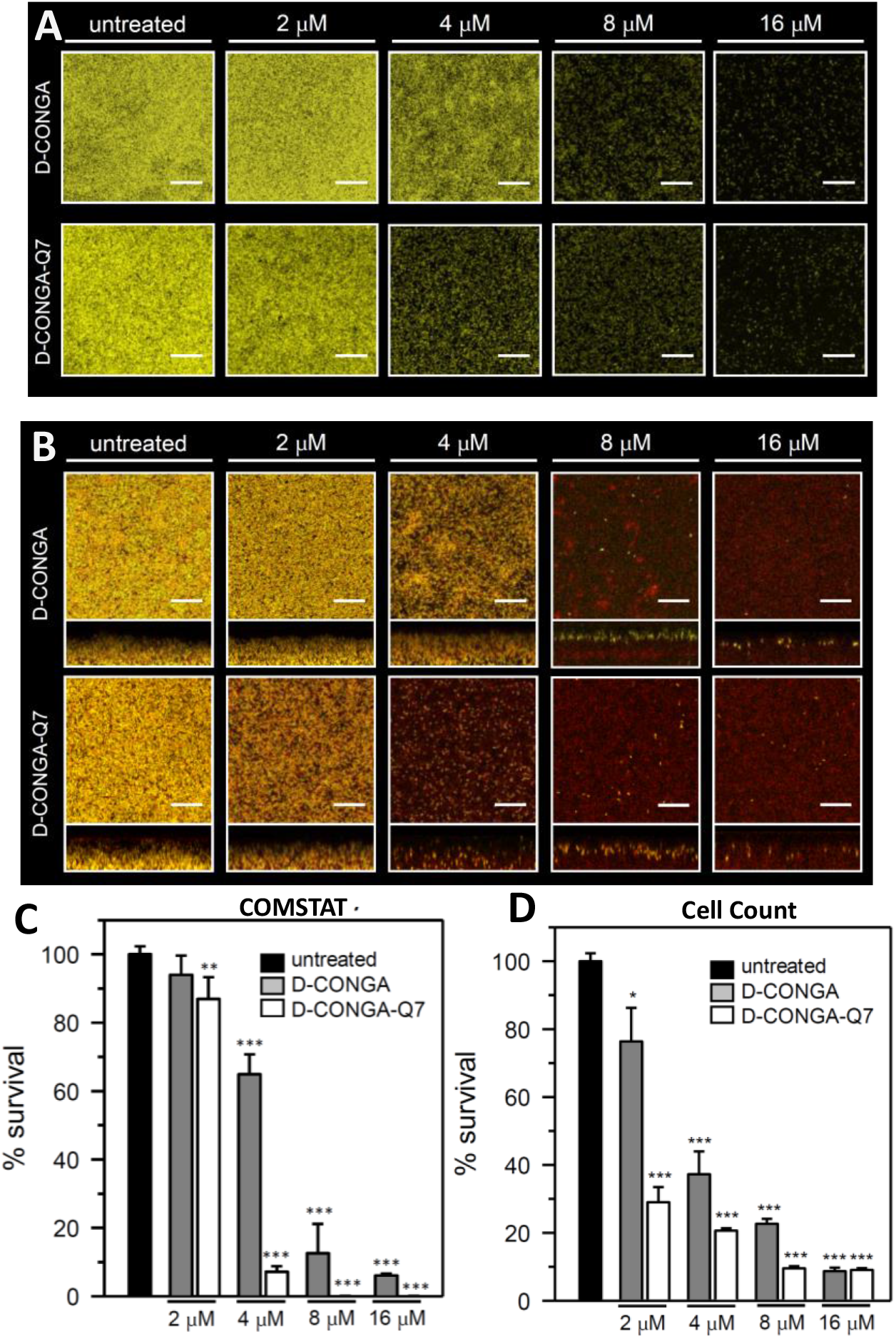
Antibiofilm activity of D-CONGA and D-CONGA-Q7. Pellicle biofilms of *P. aeruginosa* PAO1 expressing the enhanced yellow fluorescent protein (EYFP) were cultured for 48 h in PI media supplemented with 20 μM Fe and then treated for 24 h with different concentrations of peptide. For CLSM imaging, the biofilms were counterstained with Sytox Red, which stains only the DNA released from dead cells or exposed within dead cells, and imaged by CLSM. **A.** Images show the maximum projection of viable cells (yellow) after treatment. **B.** Images depict top-down (squares) and side views (rectangles) of viable cells (yellow), dead cells (red) and extracellular DNA (red). The scale of the bars represents 20 μm. **C.** The % cell survival calculated from the CLSM images with the aid of COMSTAT software. **D.** The % cell survival calculated from CFUs after dispersing the biofilms and enumerating viable cells. *p* < 0.1 denoted by *, *p* < 0.01 by ** and *p* < 0.001 by *** relative to untreated.

In **Fig. 7**, we show a direct comparison of the antibiofilm activity of D-CONGA and D-CONGA-Q7 to the clinically used lipopeptide antibiotic, colistin. Both synthetically evolved peptides are substantially more active against *P. aeruginosa* biofilms than colistin. For example, while colistin shows no detectable activity at 2 μM, D-CONGA reduces biofilm CFU by 20% and D-CONGA-Q7 reduces biofilm CFU by 70%, at the same concentration.

**Figure 7.**
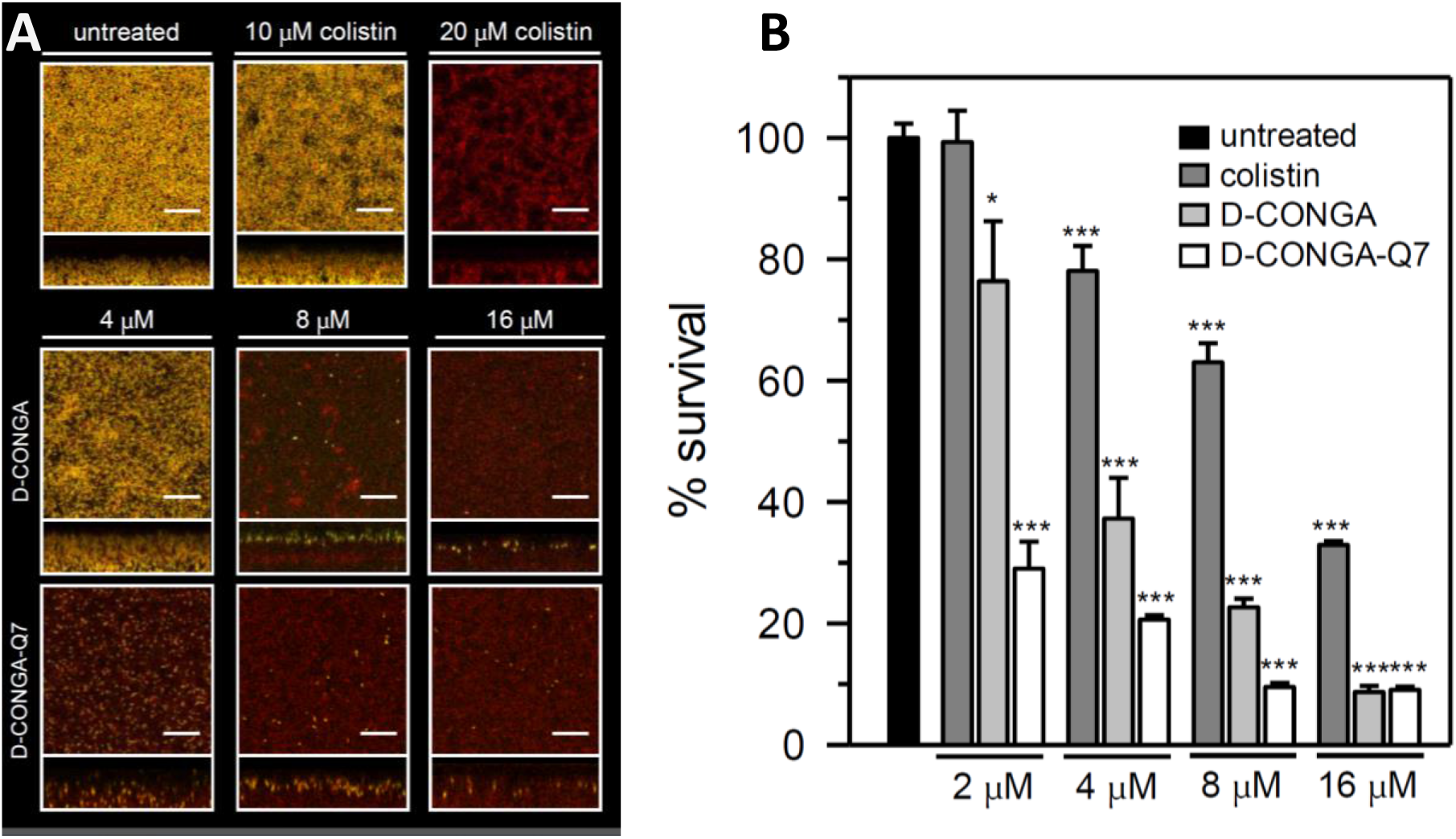
Comparison of the antibiofilm activity of D-CONGA and D-CONGA-Q7 with colistin. Pellicles of *P. aeruginosa* PAO1 expressing the EYFP were grown for 48 h in PI media supplemented with 20 μM Fe and challenged with peptides or colistin for 24 h. **A.** The biofilms were counterstained with Sytox Red, which stains only the DNA released from dead cells or exposed within dead cells, and imaged by CLSM. Images depict top-down (squares) and side views (rectangles) of viable (yellow) and dead cells and extracellular DNA (red). The scale of the bars represents 20 μm. **B**. Comparison of cell survival in pellicle biofilms after treatment with peptides or colistin by dispersing and counting viable cells. The % survival is expressed as the ratio CFU/mL (untreated) / CFU/mL (treated with peptides). *p* < 0.1 denoted by *,*p* < 0.01 by ** and *p* < 0.001 by *** relative to untreated.

#### *In vivo* antibacterial activity of D-CONGA-Q7 in a murine wound model

D-CONGA-Q7 was evaluated in a murine wound model that we used previously to characterize D-CONGA^5^. In this model, a deep surgical punch wound, created on the dorsal surface of healthy adult CD1 mice is stabilized with a silicone ring to prevent healing by contraction, and then, covered with a Tegaderm dressing. The wound bed, with a volume of ~20 μL, is immediately infected with 1×10^5^ cfu of luciferase-producing *P. aeruginosa* or MRSA. Within hours the wound bed is purulent and has a high bacterial burden^5, 15^, creating a challenging cell- and protein-rich environment to test AMP activity. Wounds are treated with 75 μg of D-CONGA-Q7 in 20 μL of water with 0.025% acetic acid every 8 hours for the first 5 days. The Tegaderm dressing is removed on day 3 post-infection and fixed in 2.5% glutaraldehyde for scanning electron microscopy (SEM) analysis of wound biofilms. The use of luminescent bacteria enables daily monitoring of bacterial burden in the wounds of each individual animal, as shown by the images in **Fig. 8A** and **8B**. Integrating the luminescence across the wound bed each day allows wound bacterial burdens to be monitored over time. In these healthy adult mice, innate immunity begins to clear the infections by day 4. Thus, the relevant measure of peptide effect is the reduction in bacterial burden across the peak of infection on Days 1-3. Compared to the activity of D-CONGA^5^ the activity of D-CONGA-Q7 is equivalent, with ≥4 logs of maximum reduction of MRSA burden and ~2 logs of reduction of *P. aeruginosa* burden, **Fig. 8C&8D**. Other wound criterion scores, shown in Supplemental **Fig. S1**, demonstrate that wound appearance and healing are similar in controls and peptide-treated samples, showing that the peptide is not causing tissue damage, despite its high local concentration.

Preliminary data had shown that once per day treatments in a simple aqueous vehicle was ineffective against wound infections in this model, see Supplemental **Fig. S2.** Three times per day is required for efficacy in simple media, **Fig. 8**. Therefore, we also tested the effectiveness of D-CONGA-Q7 formulated in aqueous suspensions of carboxymethyl cellulose and Xanthan gum, which are anionic carbohydrate polymers that create highly viscous solutions at ~1% w/v concentration. We tested D-CONGA-Q7 in these viscous solutions, applied to the wound only once per day, Supplemental **Fig. S3**, and show that this simple formulation once per day was effective at reducing bacterial burdens. Clinically useful formulations of these peptides may be straightforward to develop.

**Figure 8.**
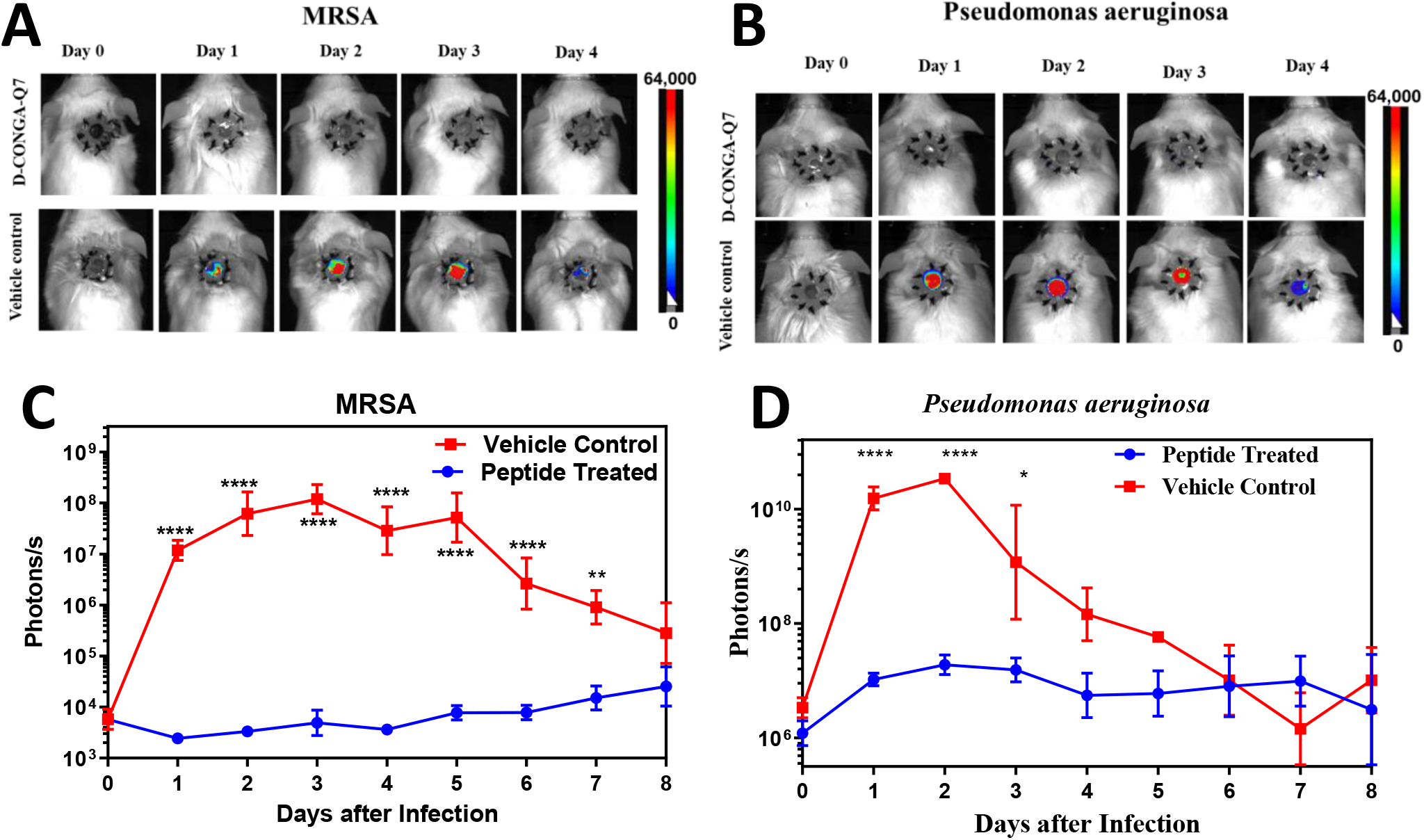
Animal model of deep surgery wound infection. Circular, dorsal puncture wounds were surgically created in healthy, adult CD1 mice, stabilized with a sutured silicon ring, and covered with Tegaderm dressing to better mimic infection and wound healing in humans. Wounds were infected with luminescent *P. aeruginosa* or luminescent MRSA and were treated with D-CONGA-Q7 peptide or vehicle control every 8 h until Day 4. An IVIS whole animal imager was used to measure luminescence in all animals once per day after infection. **A, B.** Example daily images of mice infected with luminescent *P. aeruginosa* or MRSA are shown, treated with PBS control or D-CONGA-Q7 **C, D.** Total integrated radiance from the wound bed was measured daily. Statistical analysis is described below in Methods. Significance of the difference between peptide and control denoted by asterisks.

#### D-CONGA-Q7 activity against wound biofilms in vivo

The Tegaderm dressing used in the murine wound model is an excellent substrate for bacterial adhesion and biofilm formation^5, 15^. In the wound experiment, the dressing is removed on Day 3 post-infection and fixed, followed by electron microscopy analysis. We imaged 3-4 randomly selected areas of each Tegaderm sample at two magnifications each. Each image is scored for the presence/absence of bacterial cells, either rod-shaped *P. aeruginosa*, ~1 μm in diameter, or spherical MRSA cocci, ~0.9 μm in diameter. Images are also scored for biofilm-like structures defined by having multiple bacteria embedded in an obvious three-dimensional matrix. These distinctions are shown by representative images of the fixed Tegaderm dressings in **Fig. 9A** and **9B**. In vehicle control samples, we observed that *P. aeruginosa* frequently forms an open, three-dimensional matrix in which many cells are entangled in μm-long fibers. We observed that MRSA also forms a matrix in which cells are close-packed in strands that comprise a large-scale mesh. Every SEM image (68/68) from vehicle control samples of the two bacteria contained obvious visible bacteria, and most of these images contained ≥ 50 individual cells. In samples from peptide-treated animals, 57 of 68 (84%) of images had no visible bacteria. Less than 50 bacteria were observed in each of the remaining 11 of 68 (16%) of total images. Apparent biofilms were observed in 43/68 (63%) of vehicle treated samples, while no biofilm was observed in any of the 68 images from D-CONGA-Q7-treated animals. We performed a statistical analysis of observed bacteria and biofilms using the binomial equation. Results are shown in **Fig. 9C.** P-values for all comparisons of treated vs vehicle were less than 0.001.

**Figure 9.**
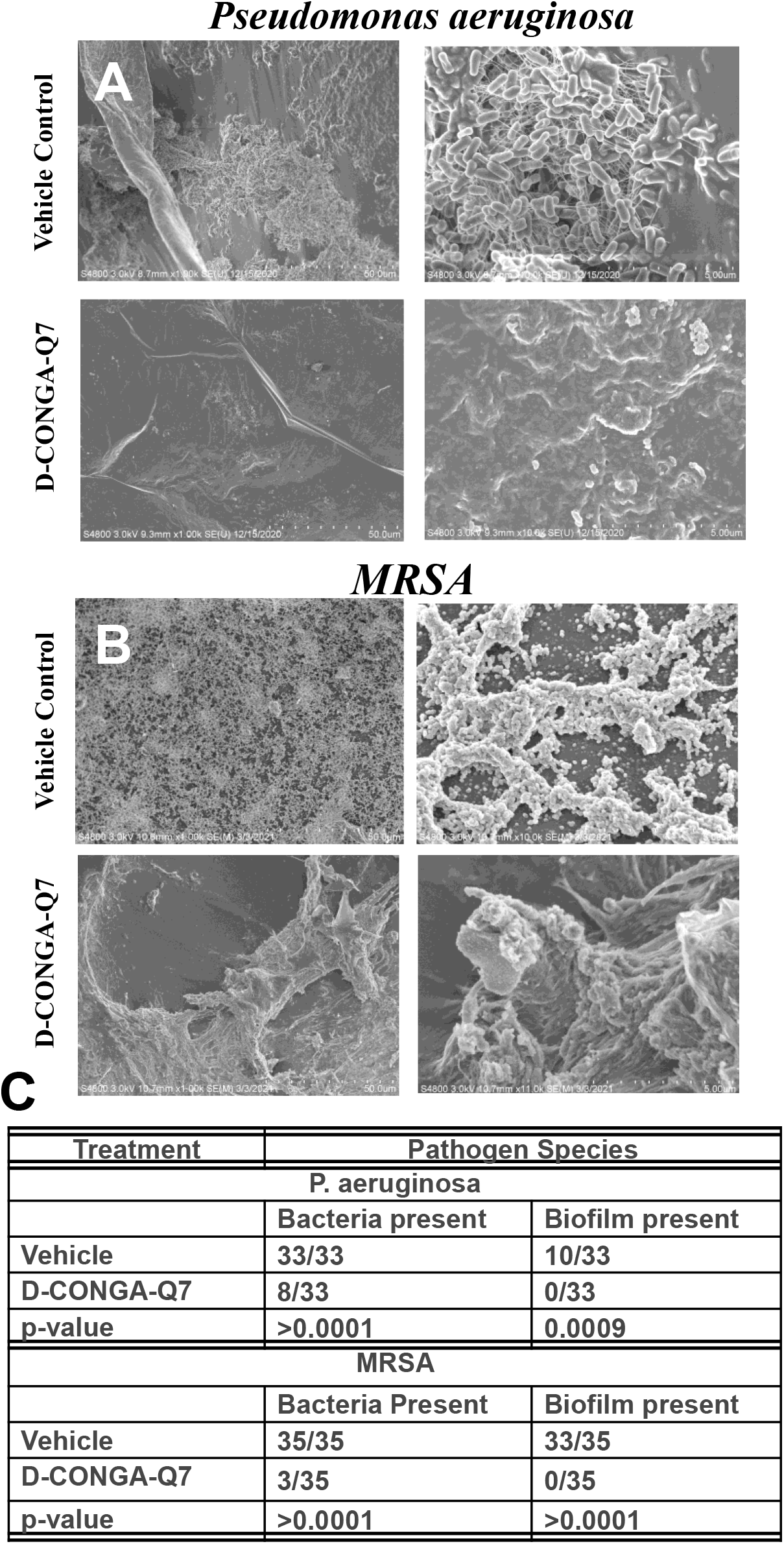
Wound biofilm reduction. **A,B.** Representative scanning electron microscopy images of glutaraldehyde-fixed Tegaderm dressing from *P.aeruginosa* infected mouse wound. The dressings removed from experimental animals on Day 3 are presented here in 1K and 10K magnifications. Scale bars are 5 μm in all images. The top two images show vehicle controls with abundant biofilms and rod-like *P.aeruginosa* in all samples. The bottom two images point on the tegaderm of wound treated 3 times a day with D-CONGA-Q7. They have few individual bacteria. No biofilm is observed, and only the tegaderm adhesive is visible. **C.** Counts of bacteria and biofilms observed in all random images of the Tegaderm dressing. P-values are determined from 2×2 contingency tables using Fisher’s exact test.

## Discussion

The overarching clinical challenge of our continued evolution of this lineage of novel AMPs is to protect and treat wounds, especially chronic wounds infected by biofilm-forming, drug-resistant bacteria in the very complex protein and cell-rich environment of an infected wound^5^. For example, current standards of care for diabetic ulcers and other chronic wounds are not effective at preventing the high rate of serious complications, especially chronic infections^26–29^. These common treatment failures are costly and negatively impact quality of life. Ultimately, complications from chronic foot wounds lead to a cumulative probability of amputation in about 16% of US diabetics^30–31^. Effective treatment for drug-resistant, infected wounds would greatly reduce the burden on both patients and healthcare providers.

Clinical use of AMPs requires that impediments be circumvented and relevant antibacterial activity be optimized. Despite decades of research into antimicrobial peptides, and thousands of distinct sequences described^4^, there is a striking absence of useful or quantitative sequence-structure-activity rules (QSAR) for any of the activities or impediments, making simultaneous parallel optimization a hopeless undertaking. Even machine learning approaches have not yet demonstrated the ability to predict clinically useful AMPs. One reason for the absence of useful QSAR is that membranes are two-dimensional fluids and peptides act on them as a dynamic ensemble of heterogeneous structures^8^. AMPs do not self-assemble into specific unique pore structures, and do not act by forming explicit pores in membranes^32–33^. Instead, AMPs accumulate massively on bacterial membranes and destabilize the membrane lipid packing by a saturation-dependent physical-chemical process that is dependent on “interfacial activity”^8^. The dependence of AMP activity on non-specific, dynamic and heterogeneous structures inhibits rational optimization. The observed properties of these ensembles are sensitive to many experimental details, leading us to suggest that they are best described by the concept of a “mechanistic landscape”^34^ rather than by a defined set of structure-function rules.

To discover new antimicrobial peptides in the absence of useful QSAR, one must resort to trial and error. In this spirit, we have been optimizing the lineage of AMPs discussed here using multiple iterative generations of library screening and rational hypothesis testing. Here, we describe the properties of the most recent generation of rational variants of the evolved, host-compatible AMP D-CONGA.

### Antimicrobial activity

The most dramatic loss of antimicrobial activity was observed for the cyclized peptides. These cyclic variants were designed to promote β-sheet or β-hairpin structure such as that found in the Θ-defensins^35^. The results thus reveal the importance of linear amphipathicity and lack of secondary structure for this peptide lineage. Further, the double arginines on both N- and C-termini are the most strongly selected feature of all previous screens^5, 11^ suggesting a critical role for this element of linear amphipathicity which is lost in the cyclized variants. Deletion of the alanine at position 4, in the N-terminal polar RRW**<u>A</u>**RR sequence, had essentially no effect on antimicrobial activity or cytotoxicity. On the other hand, deletion of the alanine at position 10, which is in the C-terminal non-polar segment LAF**<u>A</u>**FRR caused a significant reduction in antimicrobial activity, especially against *S. aureus*. This observation agrees with our hypothesis that the net length and hydrophobicity of the C-terminal nonpolar segment is critical for activity. Other observed changes are inexplicable. For example, we do not know why several seemingly unrelated variants, -βA7, ΔR13 and -LR5,LA10, lost all activity against *S. aureus*, while maintaining activity against the two Gram-negative pathogens.

### Cytotoxicity

Cytotoxicity results from the study of D-CONGA variants support our hypothesis that reducing peptide secondary structure propensity reduces cytotoxicity but does not necessarily reduce antimicrobial activity, at least against Gram negative bacteria. The two variants that we expect to be most disrupted, - βA7 and -LR5,LA10, have no detectible cytotoxicity, with EC_50_ estimated to be greater than 1 mM. There is some precedent for this idea. For example, the α-helical bee venom peptide melittin has good antibiotic activity but is also extremely cytotoxic, while a melittin diastereomer, containing the same amino acid composition in mixed L- and D-form has similarly potent antimicrobial activity, but greatly reduced cytotoxic activity^33^. This principle may become an important principle in future antimicrobial peptide design.

### Resistance avoidance

When selecting the two sets of clinical isolates to study in this work we focused on *K. pneumoniae*, a Gram-negative pathogen for which drug resistance is a serious and growing public health threat^16^. Among the clinical isolates tested, some were from pulmonary reservoirs in cystic fibrosis patients^36–38^. These strains, subjected to years of antibiotic treatment, frequently activate multiple parallel mechanisms of pan drug resistance^38^. Other isolates are from outbreaks of nosocomial infections and are known to be resistant to colistin.

We had previously passaged *P. aeruginosa* against D-CONGA over many generations and found no change in susceptibility^5^, indicating that resistance is slow to arise in *P. aeruginosa* against this peptide. In parallel experiments, resistance of *P. aeruginosa* to multiple conventional antibiotics grew rapidly over just a few passages^5^. Therefore, because Gram-negative *K. pneumoniae*, like *P. aeruginosa*, becomes resistant to some AMPs by mechanisms that affect the net charge and architecture of the outer membrane lipid A and LPS^39–42^, we did not expect resistance to D-CONGA in *K. pneumoniae*. Yet, against the *K. pneumoniae* isolates, we observed that more than half were resistant to D-CONGA^5^ whereas 28 of 29 *K. pneumoniae* isolates were susceptible to D-CONGA-Q7. This superior activity was observed despite not specifically selecting for resistance avoidance in this round of variations and testing. The mechanism of resistance avoidance is currently unknown. Further studies will be required to understand the mechanistic basis for the resistance avoidance of D-CONGA-Q7 against *K. pneumoniae* isolates. These insights will be particularly important for the advancement of resistance-avoiding AMPs into the clinic.

## Acknowledgments

AS and MR were supported by NIH grants AI125529 and AI169344. WW, LM, and JK were supported by NIH grant AI154284. JK was supported by the Louisiana Board of Regents Endowed Chairs for Eminent Scholars program, Cystic Fibrosis Foundation grant KOLLS21I0, as well as by PHS grant R35HL139930 and NIAID/NIH award R01AI114697. We thank Alan Grossfield for naming D-CONGA in Santa Fe.

Supporting Information Available free of charge: Supplemental data plots containing auxiliary information.

## Methods

### Synthesis of the peptide variants

All peptides used in this study were synthesized using solid-phase FMOC chemistry and purified to >95% using high-performance liquid chromatography (HPLC) either in the laboratory or by Bio-synthesis Inc. In all cases, peptide identity was confirmed through MALDI mass spectrometry. Unless otherwise stated, all solutions were prepared by dissolving lyophilized peptide or antibiotic powders in 0.025% (v/v) acetic acid in water. Peptide concentrations were determined by optical absorbance at 280 nm.

### Bacterial strains and growth conditions

*E. coli* (ATCC 25922), *S. aureus* (ATCC 25923), and *K. pneumoniae subsp. pneumoniae* (ATCC 13883), were used for the comparison of MIC values in this study. Subcultures, prepared by inoculating 25 mL of fresh tryptic soy broth (TSB) with 200 μL of an overnight culture, were grown to log phase (OD600 = 0.3-0.6), after which cell counts were determined by measuring the OD600 (1.0 = 5×10^8^ CFU/mL for *E. coli*, 4×10^8^ CFU/mL for *K. pneumoniae*, 4×10^8^ CFU/mL for *P. aeruginosa*, 1.5×10^8^ CFU/mL for *S. aureus*).

Bacterial cells were diluted to appropriate concentrations in TSB. We used the following clinical bacterial isolates of *Klebsiella pneumoniae* in this study: Strains KP6, KP46, NDM-1, KP396, and KP398 have been previously described^43^. ST58 strains C2, C3, and I2 are also described^41^. The alternate KP isolate is ATCC 33495. MRSA, multidrug resistant *Staphylococcus aureus* strain, is SAP400, a USA400 strain of community acquired MRSA. AB, *Acinetobacter baumannii*, is a pan drug resistant (PDR) strain isolated in the Tulane Hospital in 2015. The second independent set of *K. pneumoniae* isolates are previously described^20–21^.

The susceptibility of biofilm-embedded *P. aeruginosa* cells to the antimicrobial peptides was studied as reported previously^23^ using a *P. aeruginosa* PAO1 strain expressing enhanced yellow fluorescent protein (EYFP)^22^. The EYFP-expressing *P. aeruginosa* strain was routinely grown in *Pseudomonas* Isolation (PI) media (20 g/L peptone, 0.3 g/L, MgCl2·6H2O, 10 g/L, K2SO4, 25 mg/L irgasan, and 20 mL/L glycerol, pH 7.0). Pellicle biofilms were treated in AB minimal media^44^ with trace metals [0.15 μM (NH_4_)_2_MoO_4_, 3 μM CuSO_4_, 2 μM Co(NO_3_)_2_, 9.4 μM Na_2_B_4_O_7_, and 7.6 μM ZnSO_4_], 3 mM glucose and 15 μM Fe. The media used for the treatment included a final concentration of 0.0025% acetic acid. Iron supplementation was carried out by addition of a small volume of filter-sterilized 10 mM (NH_4_)_2_Fe(SO_4_)_2_ (pH ~ 2.0) solution. The antibiotic colistin was used at concentrations equivalent to 12.5× and 25× the reported MIC = 1 μg/mL = 0.79 μM^45^. Antimicrobial peptides stock solutions in 0.025% acetic acid were freshly prepared, stored at 4 °C, and diluted in the AB challenge media used to treat biofilms at the concentrations specified in the figure captions.

### Human red blood cells

Human O+ erythrocytes were obtained from Interstate Blood Bank, Inc. Red blood cells were subjected to four or more cycles of centrifugation at 1000xg with resuspension in fresh PBS. Following the final wash step, the supernatant was clear and colorless. RBC concentration was determined using a standard hemocytometer.

### Broth dilution assay

Antimicrobial peptides and conventional antibiotics were prepared at 5-times the final concentration needed in 0.025% acetic acid. The antibiotics were serially diluted by a factor of 2:3 horizontally across 96-well plates from Corning, 25 μL per well. One column was reserved for controls. For the assays performed in the presence of RBC, type O+ human RBCs at 0 or 2.5×10^9^ cells/mL were added in 50 μL aliquots to all wells. Following a 30-minute incubation, 50 μL of TSB, inoculated with 5 × 10^5^ CFU/mL, was added to all wells, and plates were incubated overnight at 37 °C. Following overnight incubation at 37° C, the OD600 was measured (values of less than 0.1 were considered sterilized). To assess bacterial growth in the assays with RBC, the OD600 was measured after a second day inoculation with 10 μL of solution from the original plate added to 100 μL of sterile TSB followed by overnight incubation.

#### Statistical analysis of broth dilution

Experiments were repeated 4-8 times independently, with pairs of replicate measurements on each 96-well plate. After overnight growth, optical densities were measured at 660 nm and the sterilized wells were identified by having O.D. <0.08. Non-sterilized wells were nearly opaque with OD > 0.6. For each measurement, we recorded the largest dilution that sterilized the bacteria. Statistics are done on dilution numbers, rather than calculated concentrations, because only the former have normal distributions. Mean MIC values, expressed as concentration, were determined for each microbe/peptide combination by determining mean and SE of the sterilizing dilution numbers (d) and then converting the mean d to concentration by MIC(μM) = 75/((2/3)^d) where 50 is the starting concentration in μM and the first well is considered dilution 1. In all broth dilution experiments, SE of MIC are about 0.5 dilutions, which corresponds to SE/MIC ratios of 0.2 (20%) for all measurements.

### *Hemolysis* assay

Peptide was serially diluted in PBS starting at a concentration of 100 μM. The final volume of peptide in each well was 50 μL. To each well, 50 μL of RBCs in PBS at 2×10^8^ cells/mL was added. As a positive lysis control, 1% triton was used. The mixtures were incubated at 37°C for 1 hour, after which they were centrifuged at 1000g for 5 minutes. After centrifugation, 10 μL of supernatant was transferred to 90 μL of DI H2O in a fresh 96-well plate. The absorbance of released hemoglobin at 410 nm was recorded, and the fractional hemolysis was calculated based on the 100% and 0% lysis controls. At least 3 independent experiments were performed. Results shown are averages.

### SYTOX Green cytotoxicity assay

WI-38 human fibroblast cells were grown to confluency in T-75 flasks in complete DMEM (10% FBS). The day before cytotoxicity experiments, the 10,000 cells/well cells were plated in a 96-well tissue-culture plate. The next day, in a separate 96-well plate, peptides were serially diluted in serum-free DMEM with 0.1% SYTOX green starting at concentrations of 100 μM and 50 μM, followed by 2:3 serial dilutions horizontally across the plate. To perform the cytotoxicity assay, media was removed from the wells and replaced with the peptide solutions. No peptide and 20 μM Melp5, a highly lytic peptide, were used as negative and positive control, respectively. The plate was read for SYTOX fluorescence every 5 minutes for an hour with an excitation wavelength of 504 nm and an emission wavelength of 523 nm. Percent cytotoxicity was calculated at 60 min using the 100% and 0% lysis controls. At least three independent experiments were averaged.

### Biofilm assays

These experiments were carried out as described previously^22–23^. In brief: Starter cultures (5 mL PI media supplemented with 10 μM Fe) were grown (14 h) from a single colony shaking (220 rpm) at 37 °C. To grow the pellicle biofilms, starter cultures were diluted to OD600 = 0.001 in 4 mL PI media supplemented with 20 μM Fe, placed in 35 x 10 mm petri dishes and incubated statically at 30 °C for 48 h. The pellicles were harvested using circular (1.5 cm diameter) glass coverslips by gently allowing the surface of a coverslip to contact the biofilm at the air-liquid interface. The pellicle-adhered coverslip was washed in PBS and then deposited on top of 1.5 mL AB challenge media in a well of a 12-well microplate, with the pellicle exposed to the challenge media and incubated statically at 30 °C for 24 h. The biofilms were then washed by transferring the coverslip-adhered pellicles (biofilm facing down) into 35 x 10 mm petri dishes containing 4 mL PBS, and incubating for 5 min. To release cells from the biofilm and break the extracellular matrix, the coverslip-adhered pellicles were placed in 50 mL conical tubes containing a 2 mL suspension of zirconia beads (0.1 mm diameter, BioSpec Products), 10 mL PBS, 0.2 μg/mL alginate lyase and 0.2 μg/mL DNase, incubated at room temperature for 15 min, and then vortexed for 4 min. After sedimentation of the zirconia beads, a 100 μL aliquot was used for serial dilution and plating on PIA plates for enumeration of viable cells (CFU/mL).

#### Confocal laser scanning microscopy of biofilms

These experiments were conducted as reported previously (**1**). In brief: Pellicles were washed in PBS and then stained with SYTOX Red by placing the coverslip-adhered pellicles in 1 mL of PBS containing 2.5 nM of the fluorescent dye for 15 min. Excess dye was washed with PBS, the coverslip was mounted on a glass slide using 5 μL SlowFade (Invitrogen Life Technologies) and the edges sealed with fingernail polish. The biofilms were imaged with the aid of a Leica TCS SP8 confocal microscope (Leica Microsystems, Germany) using a HC PL apo CS2 63×/1.4 oil objective. For detecting the EYFP fluorescence the laser line was set at 506 nm and the emission range to 520-610 nm. Sytox Red fluorescence was detected with excitation at 631 nm and emission range 637-779 nm. Image stacks were acquired with a *z*-step size of 0.3 μm at randomly chosen positions and the Leica Application Suite X (LAS-X) software was used for image stack processing. Quantitative analysis was performed by determination of pellicle biomass using COMSTAT^24–25^ and the Otsu method of automatic thresholding^46^.

#### Statistical analysis of biofilm treatment

One-way ANOVA followed by Tukey’s multiple *post hoc* test was used to determine the statistical significance between the means and standard deviation of untreated *vs* treated with antimicrobial agents, with the aid of SigmaPlot (Systat Software, Inc. CA).

### Wound infection model

All animal studies strictly adhered to protocol 131, which was approved by Tulane University Institutional Animal Care and Use Committee. Female CD1 mice at 8-12 weeks of age were anesthetized via intraperitoneal injection of ketamine and xylazine at doses of 90 mg/kg and 10 mg/kg, respectively. Their dorsal surface was depilated using an electric razor and scrubbed with a chlorhexidine solution. A full thickness biopsy wound was generated using a 5 mm biopsy punch (Integra). To function as a splint for the wound, a silicon (Invitrogen) ring 0.5 mm thick with an outer diameter of 10mm and a hole with a 5mm diameter was placed over the wound and held to the skin with a surgical adhesive. The entire silicon ring was then covered with Tegaderm (3M), and further adhered using 4-0 braided silk interrupted sutures (Ethicon). Mice were given 0.05 mg/kg buprenorphine immediately following surgery as well as daily for the next two days to alleviate pain from the procedure. Wound beds were infected by penetrating the Tegaderm with an insulin syringe and injecting 1×10^4^ colony forming units (CFUs) of *P. aeruginosa* (PAO1) or MRSA suspended in 10 μL sterile PBS directly onto the wound bed. All bacteria used were pelleted during early exponential growth phase prior to infection. Four hours after infection mice were topically treated with 75 μg D-CONGA-Q7 in 0.025% acetic acid or vehicle only in a 20μL volume, by penetrating the Tegaderm with an insulin syringe and injecting the treatment directly on the wound bed. Treatment was administered every 8 hours for the first 5 days of infection. Mice were imaged daily for two weeks using the *in vivo* imaging system (IVIS)-XMRS (PerkinElmer), and bioluminescence generated from the bacteria was quantified in values of radiance (photons/sec). Weight, activity, posture, coat condition and wound condition were monitored each day throughout the duration of the experiment to ensure the wellbeing of each mouse.

#### Statistical analysis of wound bacterial burden

Wound luminescence values were recorded once per day for each animal during the experiment using an IVIS whole animal imaging system. Total luminescence values, integrated over the entire wound area, were averaged for the four animals in each group. Treated versus control values were compared for each day using a two-sample t-test. Multiple comparisons were corrected for using the method of Bonferroni.

### Scanning electron microscopy

Tegaderm dressings were removed from wounds of each mouse on Day 3 after infection. Tegaderm were washed with PBS and attached to hydroxyapatite discs placed horizontally in 24-well microtiter plates. Following this, the tegaderm was fixed by placing the discs in 200 μl of 2.5% glutaraldehyde. The fixed samples were dehydrated using increasing concentrations of ethanol and then desiccated with CO2 critical point drying. The samples were carbon coated and subjected to scanning electron microscopy with a Hitachi S-4800 high-resolution microscope.

