## Supporting information for "Optimization of host cell-compatible, antimicrobial peptides effective against biofilms and clinical isolates of drug-resistant bacteria"

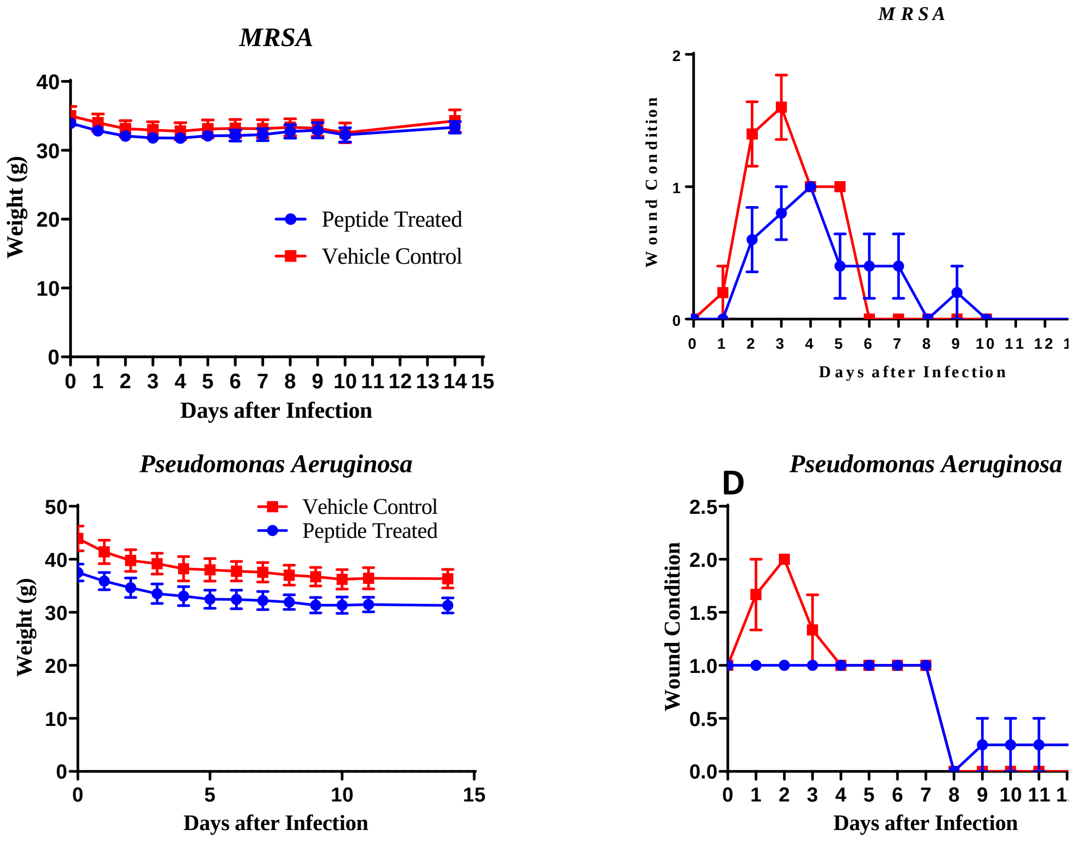


**Figure S1. Post-surgery mouse monitoring data.** The average weights (**A&C**) and wound condition score (**B&D**) of each mouse was monitored every day for the 14 days of the experiment. Uncertainties are standard errors. Each group has four mice. Mice did not demonstrate any differential weight loss based on treatment. In addition, there was no significant difference in the wound condition of mice treated with peptide versus those treated with vehicle control in *P.aeruginosa* and MRSA infected mice.

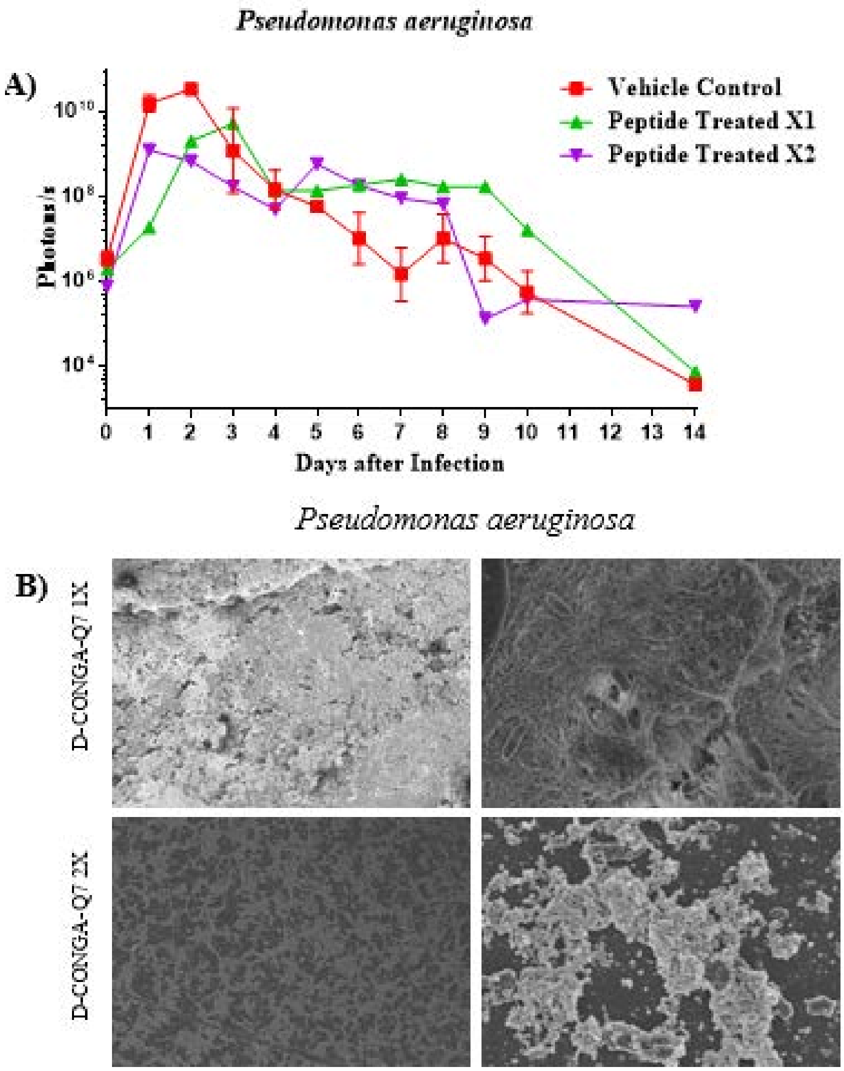


**Figure S2. Activity of D-CONGA-Q7 in wound model using 1x and 2x per day administration.** Experiments were performed as described in main text, except that treatments with 75 μg of peptide in 20 μL of buffer were administered only once or twice per day. Bacterial burdens are not reduced significantly by once or twice per day administration of D-CONGA-Q7 in PBS. Also, Tegaderm biofilm formation is not inhibited by peptide under these conditions. Results for three times per day administration are shown in **Figs. 8&9** of the main manuscript. Under this dosing regimen, bacterial burden is greatly reduced, and biofilm formation prevented.

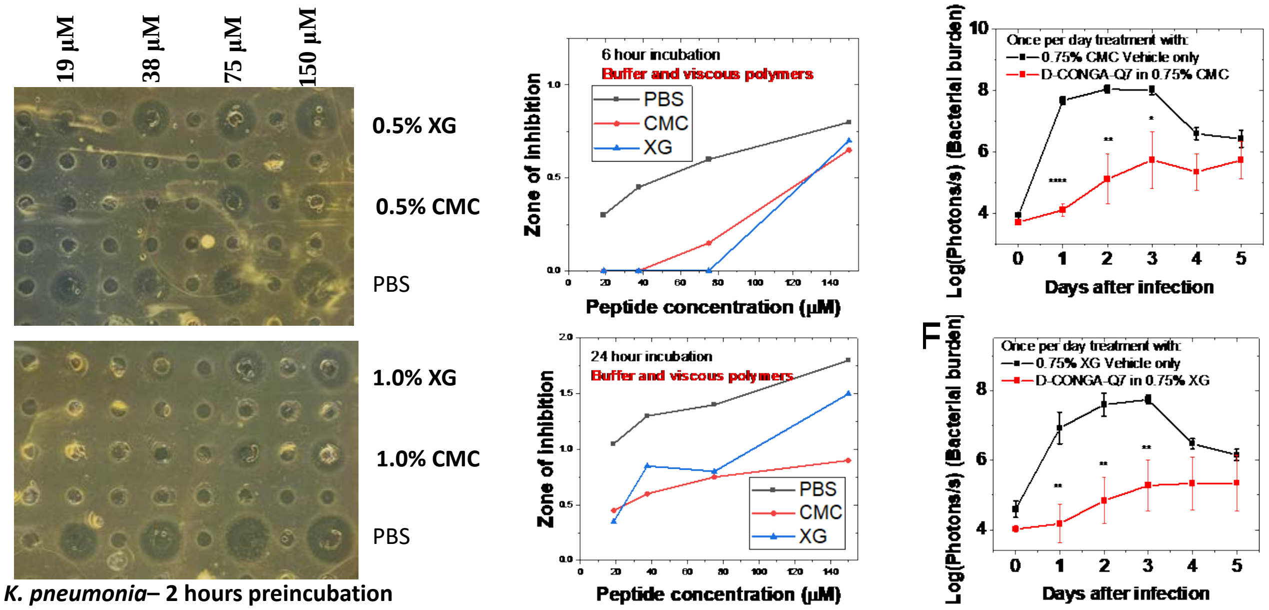


**Figure S3. Activity of D-CONGA-Q7 once per day formulated with viscous polymers**. **A-D.** *In vitro* characterization of formulated D-CONGA-Q7 using *S. aureus* in radial diffusion assays. We used 0.5% and 1% w/v suspensions of carboxymethylcellulose (CMC) or Xanthan gum (XG) to create formulations that effectively solubilized the peptide and slowed the release of peptide into agar plates for at least 24 h, compared to the vehicle, PBS. **E,F:** Wound model data for once per day administration of polymer stabilized formulations of D-CONGA-Q7 compared to vehicle PBS. Statistically significant reductions in wound bacterial burden was achieved with this once-per-day treatment.
